# Endocrinology reveals seasonal reproductive dynamics in three Malagasy fruit bats

**DOI:** 10.1101/2024.08.21.608949

**Authors:** Emily Cornelius Ruhs, Gavin K. Dehnert, Gwenddolen Kettenburg, Angelo Andrianiaina, Santino Andry, Hafaliana Christian Ranaivoson, Felix Grewe, Cara E. Brook

## Abstract

Bats (Chiroptera) are a highly diverse mammalian order with over 1,500 species, roughly 200 of which are Old World Fruit Bats (family: Pteropodidae). Many pteropodid bats, especially those residing in seasonal environments, experience seasonal reproductive cycles, including discrete periods of gestation and lactation, which underpin cyclic energetic costs. These discrete seasonal life history events have consequences for conservation management (e.g., population viability) among wild fruit bat populations; however, little is known about the seasonality of reproduction in many pteropodid bat species. Here, we aimed to elucidate the reproductive biology of three species of pteropodid bats endemic to the island of Madagascar: *Pteropus rufus, Eidolon dupreanum*, and *Rousettus madagascarensis.* We leveraged plasma samples collected in part with a longitudinal field study, from 2019 and 2020. We adapted three standard reproductive assays previously validated in humans to quantify seasonal changes in reproductive hormones for bats and applied a mixture model approach to determine hormone cutoffs indicative of pregnancy in females and mating readiness in males. As predicted, we found that pregnant females showed the highest levels of estradiol and progesterone and adult males the highest levels of testosterone. Additionally, female *P. rufus* and *R. madagascariensis* showed clear seasonality in reproduction with peaks in estradiol and progesterone in August and October, respectively. Seasonality was less clearly discernible in female *E. dupreanum* and in male testosterone data. Our results demonstrate that hormone-based reproductive classification is a powerful tool for identifying reproductive status in free-ranging bats. These physiological data refine previously reported reproductive calendars critical for demographic modeling and conservation management of threatened Pteropodidae bats.

## Introduction

Bats (order *Chiroptera*) are among the most ecologically important and diverse mammals, with over 1,500 recognized species, and roughly 200 belonging to the Old World Fruit Bat family, Pteropodidae (“flying foxes”) (Wallen 2009; Simmons and Cirranello 2026). Pteropodid bats play critical roles as pollinators and seed dispersers across tropical and subtropical ecosystems. Their survival and population stability are closely tied to seasonal patterns connected to resource availability. Yet, despite their ecological importance, many pteropodid populations are in decline due to habitat destruction and unsustainable hunting pressures (Racey 2016; Brook *et al*. 2019a; Andrianaivoarivelo *et al*. 2022; IUCN 2023), and data on their reproductive biology remain limited (Beguelini *et al*. 2021). Understanding the timing of reproduction and resulting fecundity in these species is essential for modeling population dynamics and informing effective conservation strategies (Beissinger and McCullough 2002).

Reproduction in mammals is highly regulated by endocrine signaling. In female humans and rodents, estradiol rises during follicular development, peaks around ovulation, and is followed by a progesterone-dominated luteal phase that supports pregnancy by priming the endometrium, the mucous membrane lining the uterus (Donner and Lowry 2013; Miller and Takahashi 2014). This pattern is broadly conserved in bats (Towers and Martin 1985; Santiago *et al*. 2020, 2020), though some Chiropteran species exhibit modifications of this cycle, including delayed fertilization, delayed implantation, or seasonal embryonic diapause—strategies that synchronize births with favorable environmental conditions (MacKinnon *et al*. 2003; Stukenholtz *et al*. 2018). In temperate-zone bats, estradiol typically increases during follicular development to stimulate reproductive behavior and prepare the reproductive tract, while progesterone predominates after ovulation (Crichton and Krutzsch 2000). However, the duration of the progesterone-dominated luteal phase can be extended or suppressed, depending on reproductive strategy (Crichton and Krutzsch 2000). This endocrine flexibility enables bats to synchronize birth with periods of high food availability, a critical adaptation for survival in environments with high seasonality connected to resource variability (Racey and Entwistle 2000).

For pteropodid bats, energetic demands can fluctuate dramatically across the year, particularly during reproduction, seasonal resource bottlenecks, or overlapping periods of physiological and resource demand (Andrianiaina *et al*. 2022). Estradiol and progesterone secretion can occur over extended periods, sometimes decoupled from ovulation. Many bat species exhibit a single annual birth pulse, though polyestry (i.e., two or more annual reproductive cycles; Soares *et al*. 2020) can occur for some species under favorable resource conditions (Wallen 2009; Andrianiaina *et al*. 2022). Reproductive cycles are strongly influenced by food availability and climate seasonality, and in seasonal environments like subtropical Madagascar, births are typically timed such that pups are weaned during the wet season of peak fruit availability (Arlettaz *et al*. 2001; Wright *et al*. 2005; Andrianiaina *et al*. 2022).

Despite the importance of reproductive timing to population viability for fruit bats, the endocrine mechanisms underlying these reproductive strategies remain poorly documented in most pteropodid species. Existing studies describing predictable hormonal changes during pregnancy or implantation are limited to a few species, often in captivity. Similarly, while testosterone dynamics have been described for some insectivorous and pteropodid bats (Gustafson and Shemesh 1976; Bernard 1986; Abiaezute *et al*. 2020), patterns have been shown to vary widely across bat taxa (Reeder *et al*. 2006; Duarte and Talamoni 2010). Critically, there are few validated, field-applicable tools for assessing reproductive hormones in free-ranging fruit bats. The lack of standardized methods limits our ability to characterize reproductive cycles, impeding demographic modeling and conservation planning (Ruiz-Ramoni *et al*. 2017).

This knowledge gap is particularly acute for Madagascar’s endemic fruit bats—*Pteropus rufus*, *Eidolon dupreanum*, and *Rousettus madagascariensis*. These species exhibit strong seasonal reproductive cycles, with births typically occurring between September and December each year, synchronized with the onset of the wet season (Andrianiaina *et al*. 2022). Gestation lasts approximately 3–6 months, depending on the species – with larger species gestating longer – and pups are weaned by late austral summer (Andrianiaina *et al*. 2022). Because reproductive timing in these bats is closely linked to seasonal fruiting (Ramanantsalama *et al*. 2022), even modest shifts in climate or phenology could have consequences for population dynamics. Therefore, reliable methods to determine reproductive status in wild individuals are essential for monitoring these populations.

Here, we applied a physiological approach to characterize reproductive timing in three Malagasy pteropodid bat species: *P. rufus*, *E. dupreanum*, and *R. madagascariensis*. Leveraging plasma samples from a longitudinal field study, we adapted standard reproductive hormone assays—previously validated in humans—to quantify seasonal variation in estradiol, progesterone, and testosterone for female (estradiol, progesterone) and male (testosterone) bats. We used Gaussian mixture modeling that has been previously applied to serological data to identify epidemiological states (Peel *et al*. 2013; Brook *et al*. 2019b) to classify hormone profiles into biologically meaningful reproductive states (e.g., females: nonreproductive, pregnant, lactating; males: nonreproductive juveniles, reproductively mature adults). Our goals were to add to the growing field of conservation physiology by (1) evaluating whether these hormone-based classifications can accurately be used to infer reproductive status in free-ranging bats, and (2) using these classifications to refine seasonal reproductive calendars to improve demographic models and monitoring accuracy for these specific species. By improving the ability to detect reproductive status in the field, this study provides a methodological framework that can be applied to other bat populations world-wide. These tools are directly relevant for conservation monitoring and provide an understanding of how reproduction intersects with environmental seasonality in threatened flying fox species.

## Methods

### Study site and data collection

Three species of Malagasy pteropodid bat were captured at longitudinally-monitored roosts from January 2019 to March 2020 at two main sites (e.g., Moramanga District and Ankarana National Park) in Madagascar, following methods previously described in (Brook *et al*. 2015, 2019b, 2019a) (Figure 1). Bats were classified as male or female following field protocols described in (Andrianiaina *et al*. 2022). Female bats were classified in the field into age-reproductive categories: juveniles (J), non-lactating adult (NL), pregnant adult (P), or lactating adult (L). Females were considered pregnant by abdominal palpation and lactating by presence of extended nipples or attached pups (Racey and Entwistle 2000; Mason *et al*. 2010). Note that it is more difficult to detect a pregnant bat during early pregnancy using abdominal palpation. Male bats were classified in the field as either juvenile (J) or adult (A). Plasma samples were obtained after centrifugation of blood collected into heparinized tubes from the brachial vein (<1% body mass) of captured individuals (35 male and 26 female *P. rufus*, 100 male and 144 female *E. dupreanum,* 85 male and 103 female *R. madagascariensis*), frozen in the field using liquid nitrogen, and moved to long-term storage in −80°C freezers in the capital city of Antananarivo, Madagascar. Samples were subsequently exported to the University of Chicago for hormone assays where estradiol and progesterone concentrations (n=273 females) or testosterone concentrations (n=220 males) were measured. All procedures were completed in accordance with approved animal care policies at the University of Chicago (IACUC #72675) and UC Berkeley (#AUP-2017-10-10393) and under permits from the Madagascar Ministry of Forest and the Environment (007/19, 197/19, 14/20, 182/20).

**Figure 1.**
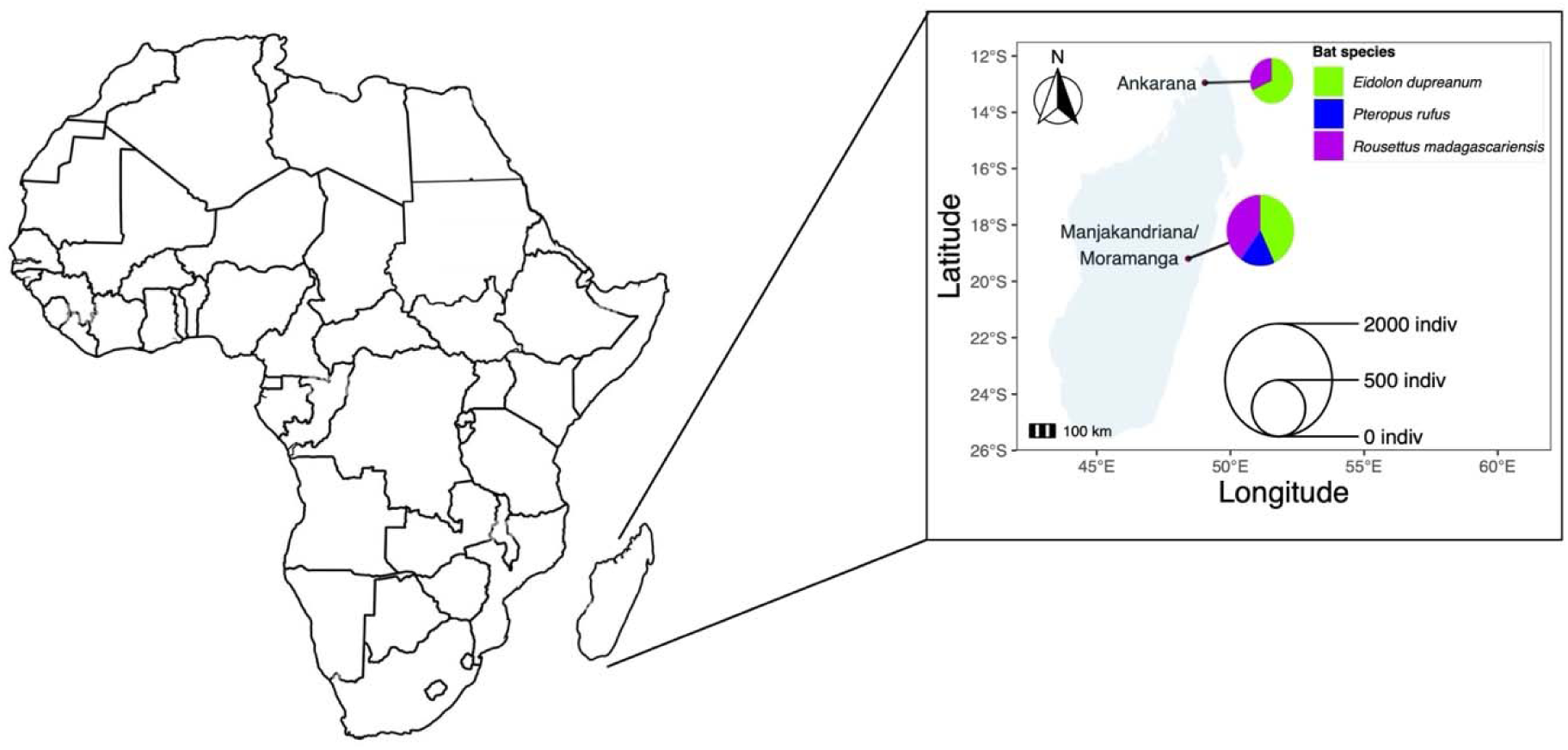
Geographic location of sampling sites used in this study. Pie charts represent the total number of each species sampled from a given site. Current inset plot is based on the sample size for the original hormone data. Plasma samples were obtained from a subset of captured individuals (144 female and 100 male *E. dupreanum,*103 female and 85 *R. madagascariensis*, and 26 female and 35 male *P. rufus,*).

### Reproductive assays

All molecular methods for determining female reproductive status were modified from methods outlined in Beguelini *et al*. (2021). Plasma concentrations of estradiol (E2) and progesterone were measured in female bat samples by two commercially available enzyme-linked immunoabsorbent assays (ELISA): estradiol E2 AccuBind ELISA Microwells (#4925-300A, Monobind, Inc. Lake Forest, CA, sensitivity 6.5pg/mL) and progesterone ELISA (#RE52231, Tecan, Inc., Männedorf, Switzerland, 0.14 ng/mL), respectively, following validation for our three species of interest (see following section). The assays were performed according to kit instructions with one modification - only 10ul of plasma was used per well. This volume was validated prior to running all samples, and subsequent concentrations were found to fall within the standard curve. Plasma concentrations of testosterone in male bat samples were measured using a commercially available ELISA: Testosterone Accubind ELISA Microwells (#3725-300, Monobind, Inc. Lake Forest, CA, sensitivity 0.576 pg/mL). The testosterone assay was performed according to kit instructions, again following validation for our three species of interest (see following section). We additionally obtained adult male *R. aegyptiacus* samples from Lubee Bat Conservancy (Gainesville, FL, USA) to help benchmark low testosterone values for *R. madagascariensis*. For each assay, final absorbance was read at 450nm and at 625nm for background correction. Samples were run in duplicate for each hormone; however, due to small plasma volumes, a handful of female samples were not analyzed for both hormones (n=6 samples only have progesterone data and no corresponding estradiol data).

### ELISA Assay Validation

We validated commercial ELISA kits for estradiol, progesterone, and testosterone for all three Malagasy pteropodids using three different techniques: ‘dilution linearity’ (whereby a predictable decline in hormone concentration is recoverable across a dilution gradient); ‘parallelism’ (whereby the slope of the dilution series for each bat is shown to parallel the assay’s standard curve [derived from humans], suggesting comparable antibody dynamics across taxa); and ‘spike and recovery’ (whereby a known concentration of each hormone is added to a bat sample with previously assayed hormone quantity, and we are able to effectively recover the spiked quantity using the specified kit). Thus, ‘dilution linearity’ evaluates the proportionality and predictability of the assay in bats, ‘parallelism’ evaluates the equivalence of hormone behavior across taxa, and ‘spike and recovery’ evaluates the assay’s accuracy and confirms a lack of inhibitory or amplifying interactions in the bat samples. For dilutional linearity and parallelism, three to six samples per species (n=6 samples *R. madagascariensis*; n=3 samples *P. rufus*; n=4 samples *E. dupreanum*) were combined, and serial dilutions 1:2, 1:4, 1:8, 1:16, 1:32, and 1:64 were performed, then compared against the standard curve. Only samples within the standard curve limit were used for dilution linearity and parallelism. For spike and recovery, samples with known concentrations of each hormone were selected, and 50uL of each individual sample was allocated to each sample tube. 50uL of a standard was additionally allocated to each sample tube for a total of 100uL (n=5 samples per species per assay). Analyses for assay validation were completed using GraphPad Prism Software 10.5.0 (GraphPad Software Incorporated, La Jolla, Ca, USA). For dilution linearity and parallelism validations, normality assumptions were evaluated using Shapiro-Wilk and Kolmogorov-Smirnov tests, linear regressions were fit to each species-hormone concentration of the data, and the randomness of the residuals from each regression was assessed with a Runs test. For spike and recovery validation, we quantified the average recovery percentage of each spike for each species-hormone concentration and the 95% confidence interval based on standard error across all n=5 replicates. Results of the validation are presented in Appendix 1, Table S1, Figures S1-S3.

### Statistical analysis

All other analyses were performed in R (Version 2022.02.2, build 492).

#### Assigning reproductive hormone concentration values

Standard curves were constructed for all of the reproductive hormones using the ‘drc’ package in R (Ritz *et al*. 2015; Ritz and Strebig 2016). Control samples of known hormone concentration and optical density (OD) were run alongside field samples to establish a standard curve. We first fit a log-logistic (LL4) model as suggested from the kit (Ritz *et al*. 2015) to the resulting data to translate OD into hormone concentration; then, projected hormone levels were averaged from duplicate reads obtained for all field samples. In some instances, for all hormones, values were either too high or too low to fit the standard curve model. OD is a measure of the transmission of light through a blocking optical filter defined as the negative log_10_ of transmission; thus, in those cases, we determined the concentration accordingly: (1) if the OD value was too high for the standard curve and concentrations were below the standard curve (meaning the animal exhibited a hormone level inconsistent with either pregnancy in females or mating readiness in males), we assigned the value of the kit lower bound sensitivity (6.5 pg/ml for estradiol, 0.14 ng/ml for progesterone, and 0.576 pg/ml for testosterone), and (2) if the OD value was too low for the standard curve and concentrations were above the standard curve (meaning the animal exhibited a hormone level consistent with either pregnancy in females or mating readiness in males), we assigned a value equal to the highest standard in the kit (estradiol = 1500 pg/ml, progesterone = 40 ng/ml, and testosterone = 12.0 ng/ml).

#### Assigning pregnancy status

To understand the correlation between estradiol and progesterone, our two measured hormones associated with pregnancy, we constructed linear models for each species in our analysis with progesterone predicting an estradiol response and, additionally, for estimated day of reproductive year predicting both progesterone and estradiol levels. Next, we conducted a PCA analysis to collapse estradiol and progesterone values into a single metric prior to determining pregnancy status, including the estimated date in the reproductive calendar for each species. For samples derived from the District of Moramanga, we assumed day 0 (parturition) to occur on September 29 for *P. rufus,* November 16 for *E. dupreanum*, and December 12 for *R. madagascariensis* and the corresponding day 365 (respectively, September 28, November 15, December 11) to correspond to late stage pregnancy in all species, following reproductive calendars reported for these species in this region in (Andrianiaina *et al*. 2022). In the Ankarana National Park sites, our field data indicated an earlier annual birth pulse for *E. dupreanum* (data were not robust enough to identify any differences in reproductive calendar for *R. madagascariensis,* and no *P. rufus* were captured in this region). Accordingly, we set the assumed parturition date to the earliest date in the dataset on which lactating female *E. dupreanum* with attached pups were observed: September 26. Because fruit bat birth pulses can be staggered across several days or weeks in a given location, in the same sampling year (2019) also at the Ankarana field site, we additionally observed four very pregnant female *E. dupreanum* a few days following this assumed birth pulse date, two each on October 1 and 2. To maintain the association between high day of year values with late pregnancy and low day of year values with parturition and the onset of lactation, we fixed these four datapoints with an assumed day of year value of 365. Doing this maintained consistency in high value metrics for estradiol, progesterone, and day of year, all associated with pregnancy and allowed for their combination in PCA analysis. The resulting PCA1 coordinates were then normalized to avoid negative values by adding 1 to each value. As male bats only had values for testosterone, we did not conduct a PCA and used raw hormone data in subsequent analyses.

Next, we conducted a cluster analysis approach to determine the cutoff values for pregnancy in females. Using the mclust() package in R, we fit gaussian mixture models to the normalized PCA values to approximate a cut-off concentration and lower and upper confidence intervals for each of the three species. The cluster analysis assumes an overlapping series of normal distributions corresponding to diverse reproductive categories; for example, nonreproductive females will exhibit estradiol and progesterone values (or the PCA1 value returned from these two combined with date) within some (lower) distribution, while pregnant females will exhibit values within some (higher) separate distribution. mClust requires at least two distinguishable clusters; therefore, we allowed the data to determine the best fit number of normally distributed clusters, ranging from 2 to 3. For females, we ran the mixture models on the normalized PCA values and allowed the number of mixture components to be set to the default (G=2:3). Female bats were classified as pregnant if their PCA1 value clustered within the uppermost distribution determined by the mixture modeling (e.g. distribution 2 or 3) with 70% certainty (Table S2; lower confidence thresholds were also set at 60% certainty and higher confidence thresholds at 85% certainty). When the cluster-based classification indicated pregnancy, but the original field designation did not, we corrected the bat’s reproductive status accordingly. There was one case where a female *R. madagascariensis* bat (ROU154) had aborted a fetus in the field. Here, the cluster approach did not detect hormone levels indicative of pregnancy; however, for our subsequent analyses, we nonetheless modeled this individual as ‘pregnant’ based on direct observation of the newly expelled fetus.

After determining reproductive status through the cluster analysis, we assessed how well these classifications aligned with our original field assignments. Specifically, we calculated the percentage of cases where the field-assigned reproductive status matched the cluster-based classification. We summarized the total number of female samples with field reproductive status data and counted how many individuals were reclassified as “P (pregnant)” by the cluster analysis. This comparison was conducted only for females.

#### Descriptive statistics for categorical status

We conducted nine analysis of variance (ANOVA) tests with hormone as the response variable and reproductive status as the predictor to determine if categorical status groups had significantly different hormone (estradiol, progesterone, and testosterone) levels. All ANOVAs were followed by a Tukey HSD posthoc analysis to determine the relative directional significance between groups.

#### Seasonal model building

We used the gam() function in the ‘mgcv’ package in R (Wood 2022) to build generalized additive models (GAMs) to understand the seasonal patterns in reproductive hormones (estradiol, progesterone, testosterone). We added the absolute minimum of the value +1 to all raw hormone data to remove any negative values and then log-10 transformed the data. As most data were derived from the central District of Moramanga region in Madagascar, GAMs in the gaussian family were fit to the data from only this region, as there are likely differences in reproductive calendar between the north (Ankarana) and central (Moramanga) regions, and seasonal representation was more complete for the Moramanga subset of the data. For all GAMs, we converted collection date of sample to a day of year metric to control for intra-annual variation as a smoothing effect, using a cyclic cubic regression spline (bs= “cc”), holding knots at k=7, as recommended by the package author (Wood 2022), and included year as a random effect (bs= “re”).

## Results

### Assigning pregnancy status

Estradiol and progesterone were significantly positively correlated for all three species of bats (Figure S4A). Estradiol/progesterone and day of reproductive year were significantly positively correlated for *E. dupreanum* and *R. madagascariensis*, but not *P. rufus* (Figure S4B and C). For the female *P. rufus*, PCA1 explained 45.0% of the variation whereas PCA2 explained 32.5% of the variation (Figure S5). For the female *E. dupreanum*, PCA1 explained 65.3% of the variation whereas PCA2 explained 25.1% of the variation. For the female *R. madagascariensis*, PCA1 explained 64.7% of the variation whereas PCA2 explained 21.5% of the variation. While PCA values have no innate biological meaning, here, we used them to summarize hormone and date of reproductive year (with day 0 set to parturition), which when combined, correlated with pregnancy status in female bats (Table S2; Figure S6).

### Reproductive analysis

Estradiol and progesterone concentrations were partially explained by reproductive status for *E. dupreanum* and *R. madagascariensis*, but not for *P. rufus*, where sample sizes were smaller (Figure 2; Table S3). Estradiol concentrations were significantly higher in pregnant *E. dupreanum* and *R. madagascariensis* compared to juveniles (p<0.001 and p=0.001, respectively), non-lactating (p<0.001 and p<0.01, respectively), and lactating (p<0.001and p<0.001, respectively) bats. Estradiol was not significantly different across any groups in *P. rufu*s (p=0.109). Progesterone was higher in *E. dupreanum* and *R. madagascariensis* compared to juvenile (p<0.001 and p<0.001, respectively), non-lactating (p<0.001 and p=0.03, respectively), and lactating (p<0.001 and p<0.001, respectively) bats. Progesterone was not significantly different across any groups in *P. rufus* (p=0.623).

**Figure 2.**
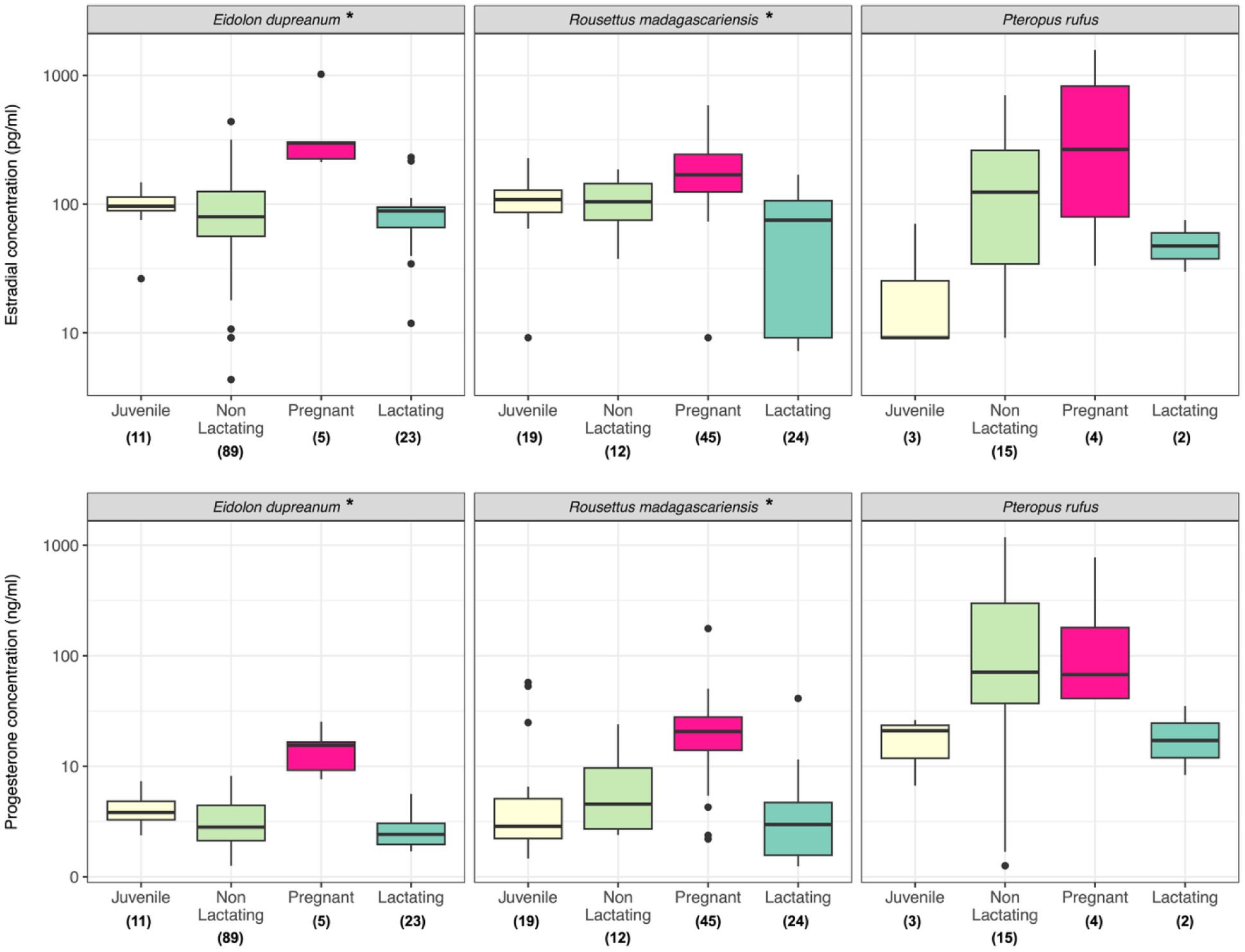
Estradiol (pg/ml) and progesterone (ng/ml) concentrations for female bats plotted by reproductive status. Samples sizes are listed in parentheses under each group. Asterisks in the header indicate a significant (p<0.05) ANOVA result. If females were determined to be pregnant via hormonal assay, catching status was switched to pregnant (‘P’) for this analysis. See methods section for more details.

Hormone analyses suggest potential misclassification of reproductive status in all three species (Table 1). The highest rate of misclassification from laboratory hormone analysis was in the non-lactating adult female group (NL), which ranged from 1.1% for *E. dupreanum*, to 63.6% for *R. madagascariensis* and 11.8% *P. rufus*. For hormone analysis, adult lactating (L) bats were rarely misclassified in the field. There were some rare instances where females were identified as pregnant in the field, and the hormone analysis did not confirm pregnancy status (n=3 *R. madagasacariensis*; n=2 *P. rufus*). For *R. madagascariensis and P. rufus*, these samples were captured during late January and July in Moramanga District, respectively.

**Table 1.** Summary statistics of female bats that were classified correctly in the field according to the reproductive analysis. If female bats had hormonal levels above the cutoff values (see main text) their “status” was reclassified as “P” (pregnant); however, if their hormonal levels were below the cutoff values, their catching status from the field was maintained. Statistics below indicate the percentage of the time that the field status matched the status assigned by the reproductive assays (e.g. proportion matched) or that they did not match (e.g. proportion mismatched).

| *Species* | *Sex* | *Catch status* | *Total count sampled* | *Count of matched* | *Proportion match* | *Proportion mismatched* |
|---|---|---|---|---|---|---|
| *Eidolon dupreanum* | Female | Female - all | 128 | 127 | 0.992 | 0.008 |
|  | Female | L | 23 | 23 | 1.0 | 0.0 |
|  | Female | NL | 90 | 89 | 0.989 | 0.011 |
|  | Female | P | 4 | 4 | 1.0 | 0.0 |
| *Pteropus rufus* | Female | Female - all | 24 | 22 | 0.917 | 0.083 |
|  | Female | L | 2 | 2 | 1.0 | 0.0 |
|  | Female | NL | 17 | 15 | 0.882 | 0.118 |
|  | Female | P | 2 | 2 | 1.0 | 0.0 |
| *Rousettus madagascariensis* | Female | Female - all | 100 | 78 | 0.780 | 0.220 |
|  | Female | L | 25 | 24 | 0.960 | 0.040 |
|  | Female | NL | 33 | 12 | 0.364 | 0.636 |
|  | Female | P | 24 | 24 | 1.00 | 0.000 |

Testosterone values were generally low for all species and showed weak or inconsistent seasonal trends, though values remained consistently low in *R. madagascariensis* regardless of age class. Adult *P. rufus* males had significantly higher testosterone concentrations from juveniles (p=0.03); however, this trend was only marginally significant for *E. dupreanum* (p=0.057) and not significant for *R. madagascariensis* (p=0.71; Figure S7; Table S3). While we attempted multiple testosterone ELISA kits to attempt to recover clearer data for *R. madagascariensis*, we observed that *Rousettus aegyptiacus* testosterone values (which we assayed for additional verification) were also quite low (0.152 ± 0.06 pg/ml); thus, it is possible the genus *Rousettus* has low circulating values of testosterone in general.

### Effect of seasonality

Both estradiol and progesterone demonstrated significant intra-annual seasonality among all three bat species (estradiol, *E. dupreanum*: p<0.001, *P. rufus*: p<0.001; progesterone, *E. dupreanum*: p<0.001, *R. madagascariensis*: p=0.002; *P. rufus*: p=0.002), except for female *R. madagascariensis* estradiol levels (p=0.84; Figure 3). Notably, GAMs to test for seasonality were fit to data from the central District of Moramanga field sites only, and the majority of samples corresponding to the gestation period for both *E. dupreanum* and *R. madagascariensis* were collected from the northern Ankarana National Park sites, excluded from seasonal analysis (Figure 3). Year was a significant random effect for *E. dupreanum* for estradiol (p=0.003) and *P. rufus* for progesterone (p=0.003), but not the other species-hormone combinations.

**Figure 3.**
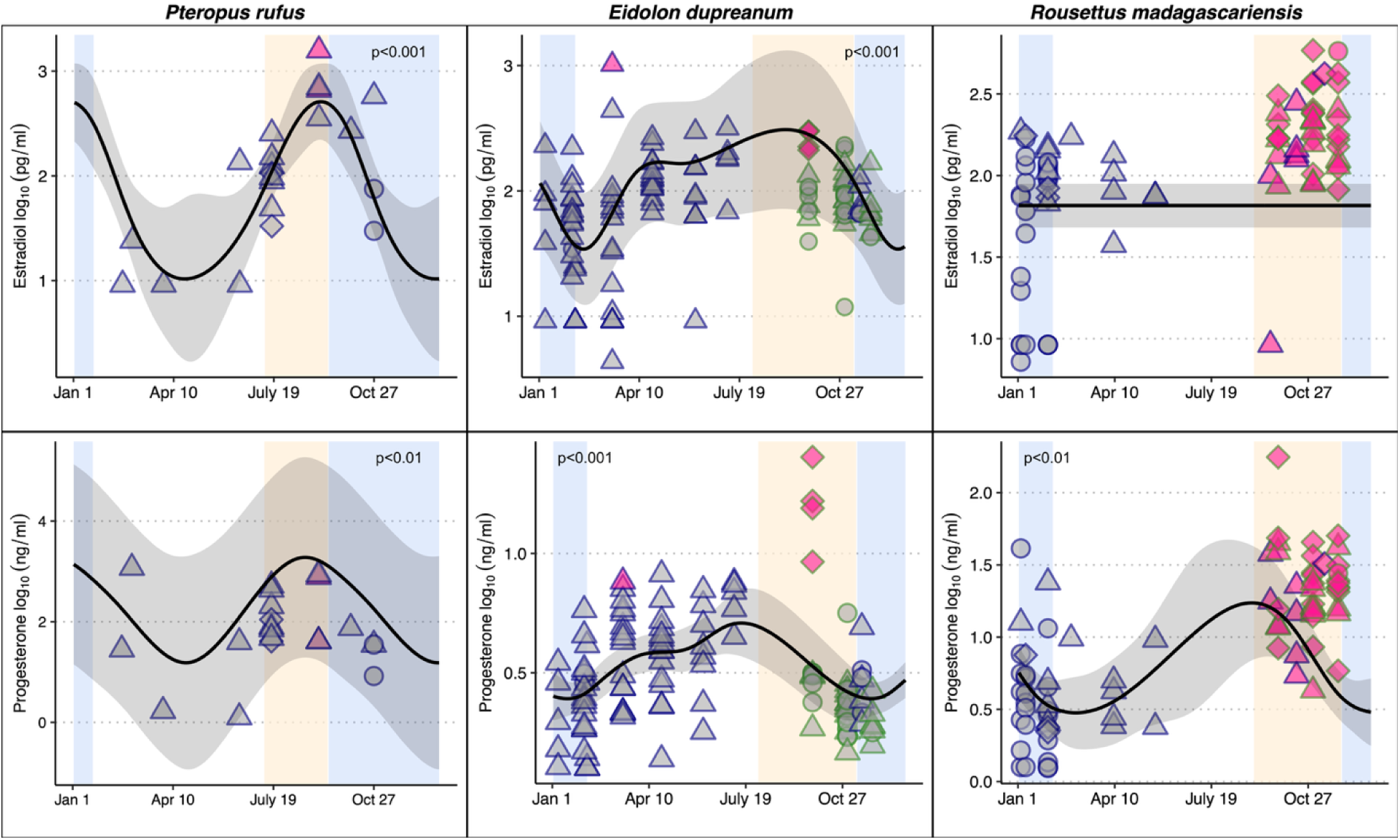
Seasonal patterns of estradiol and progesterone for all three species of bats. *P. rufus* on left, *E. dupreanum* in the middle, and *R. madagascariensis* on the right. Shape of the points indicate reproductive class by catching status (lactating = circles, non-lactating = triangles, and pregnant = diamonds), as evaluated in the field, while bats identified as pregnant vs. non-pregnant by hormone assay are colored, respectively, pink vs. gray. Points outlined in dark blue are from the District of Moramanga whereas points outlined in green are from Ankarana. The duration of gestation is highlighted in light peach and lactation in light blue for each species for the District of Moramanga region. Solid line indicates generalized additive model trends with 95% confidence intervals shaded in gray (GAMs were fit to District of Moramanga data only). All female hormones showed seasonal effects by day of the year, except for estradiol in *R. madagascariensis*.

Testosterone showed significant intra-annual seasonality only for *E. dupreanum,* with no clear seasonal structure for *P. rufus* or *R. madagascariensis* (Figure S8). These patterns may reflect lower or less synchronized male reproductive activity in this system. Field observations have suggested that *R. madagascariensis* bats may have multiple annual birth pulses at lower latitudes but only one pulse at higher latitudes (Andrianiaina *et al*. 2025), in agreement with previous work in *R. aegyptiacus* (Lučan *et al*. 2014).

We combined information from reproductive hormone assays to attempt to quantify seasonal changes in the reproductive calendar for all three Malagasy fruit bats and compare against the previously reported reproductive calendar first presented in Andrianiaina et al. 2022 (Figure 4).

**Figure 4.**
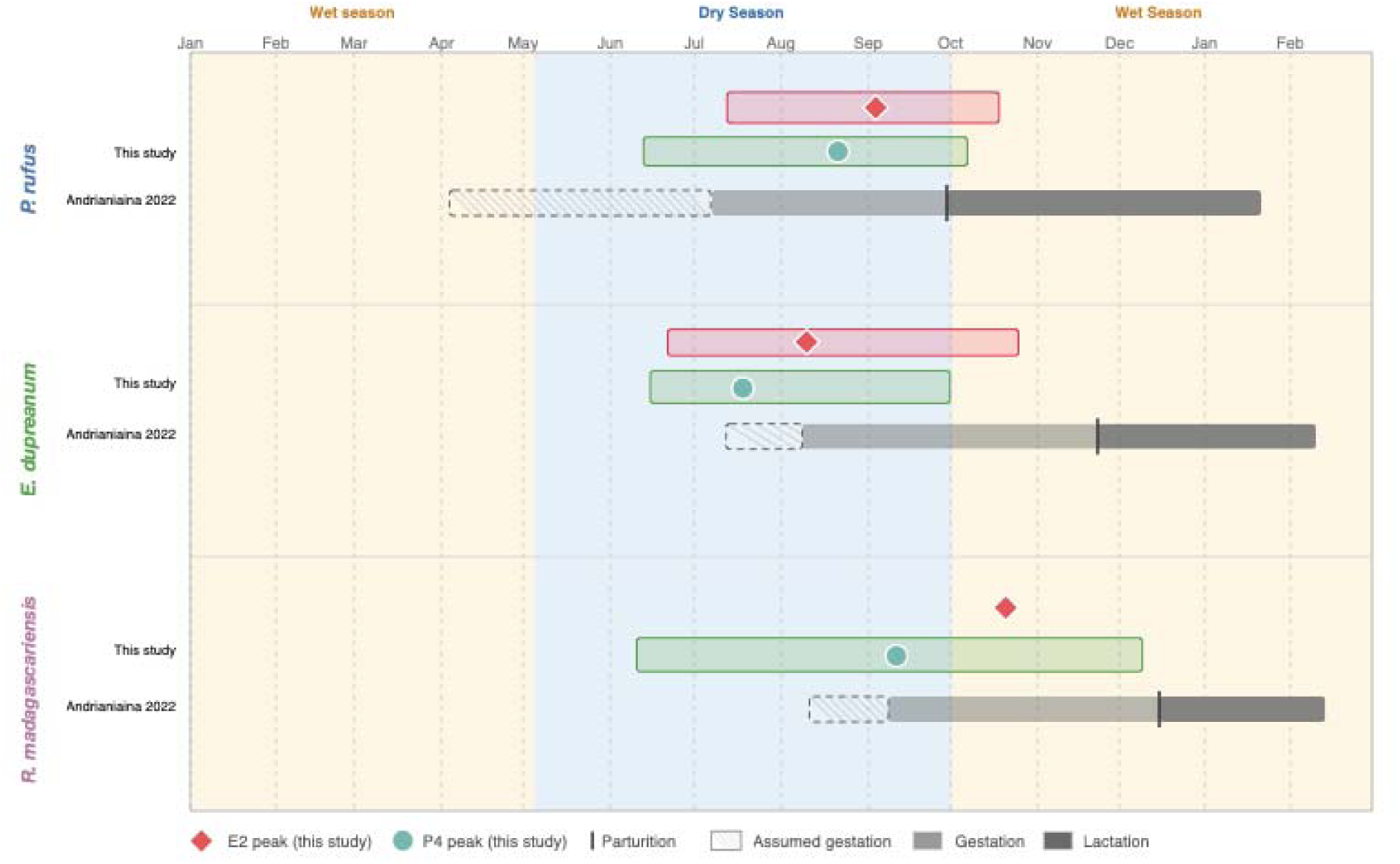
Schematic comparison of reproductive timing between hormone-derived estimates (this study) and previously-reported field-based observations for fruit bats in the District of Moramanga (Andrianiaina *et al*. 2022). Red points and bars depict estradiol measurements while green points and bars depict progesterone measurements. The points indicate the seasonal timing of the peak value for each respective hormone from our GAM analysis (Fig. 3), while the bars depict the range spanning from the midpoint of the inflection line between the GAM peak and the first trough to its left to the midpoint of the deflection line between the GAM peak and the first trough to its right. Bar position denotes timing rather than magnitude. As we observed no seasonal patterns for estradiol in *R. madagascariensis* in this analysis, we only depict the peak value and not an estimated window (e.g. with a bar).

## Discussion

Accurate classification of reproductive status for Old World Fruit Bats (Pteropodidae) has significant implications for understanding basic population demography and informing conservation efforts. Here, we characterized female and male reproductive hormones levels and developed seasonal species-specific profiles of two pregnancy-related hormones, progesterone and estradiol, using samples from a longitudinal field system in Madagascar. Our results confirmed that estradiol and progesterone can be accurately measured using commercially available assays, previously never validated for these three species. Furthermore, our study indicates that these hormones can be used as indicators of different reproductive life stages (e.g., juvenile, non-lactating, pregnant, and lactating), which can be determined without sacrificing an animal.

Progesterone and estradiol values for pregnant *E. dupreanum* and *R. madagascariensis* bats were significantly higher than those observed in all other reproductive classes (juveniles, non-lactating, and lactating female bats) (Table S3). The values for estradiol and progesterone observed in pregnant Malagasy fruit bats of all three species were similar to those previously observed during other bat species in the late stages of gestation, prior to the decline observed during the lactation period (Figure 2); however, we note that other studies reporting reproductive hormone levels in wild caught bats used destructive sampling to confirm pregnancy stage. For example, *Artibeus lituratus,* a small-sized neotropical bat, showed steady increases in both estradiol and progesterone from early to late stages of pregnancy (“P3”), with a subsequent decline after birth (Santiago *et al*. 2020). Similarly, in *Myotis nigricans*, a small insectivorous bat, serum concentrations of estradiol and progesterone in early pregnancy (“P1”) and late pregnancy (“P2”) were higher compared to the lactation stage (Beguelini *et al*. 2021). While trends in reproductive hormone values reported for other bat species mirror the trends presented here (e.g., elevated in pregnant bats), overall ranges are species-specific.

Our previous study combined field-data with values from the literature to elucidate the reproductive cycle for the same three species of bats investigated here (Andrianiaina *et al*. 2022). Previously, we estimated that all three species exhibited a single annual birth pulse in the central Moramanga District, with gestation ranging from 120-180 days and the birth peak occurring between September-December, depending on the species (Andrianiaina *et al*. 2022). The data generated here are largely in agreement with those previously reported in Andrianiaina et al., (2022)—with the exception of *E. dupreanum*, which demonstrated high female reproductive hormone levels throughout the year in our analysis (Figure 4). It is possible that *E. dupreanum* may either have consistently higher hormone levels than other species; have an earlier birth pulse than previously assumed (∼July/August instead of November; Andrianiaina *et al*. 2022); or have biannual peaks in estradiol levels, with the earlier peak corresponding to fertilization and the latter peak to gestation (Figure 3). While much is still unknown about the reproductive biology of Malagasy fruit bats, elucidating pregnancy timing to inform demographic models will allow more accurate forecasts of population trajectories, particularly in the face of climate-related threats, including changes in fruiting phenology and extreme weather events.

For female bats, both estradiol and progesterone (with the exception of estradiol for *R. madagascariensis*) showed seasonal signatures – with both hormones increasing in the mid to late dry season. Previous reports of the reproductive calendar for these bats are in line with the seasonality observed here, where gestation occurs during the dry season and the birth pulse coincides with the onset of the rainy season which corresponds to fruit abundance (Andrianiaina *et al*. 2022). Moreover, while our seasonal analysis was restricted to central Madagascar, our longitudinal sampling consisted of locations in both central and northern parts of Madagascar. Seasonal patterns are likely to shift based on latitude (Lučan *et al*. 2014), with bats from northern Madagascar likely breeding earlier (Walker *et al*. 2019; Andrianiaina *et al*. 2025). Unfortunately, very little is known about how the shift in timing of the resource-rich season might influence or have carry-over effects on the timing of bat breeding (Walker *et al*. 2019). Thus, future field studies should aim to increase sample sizes of fruit bats in the north of Madagascar to explore these latitudinal shifts in reproductive timing over multiple years and to elucidate how shifts in the resource availability influence hormonal levels and ultimately, birth pulses.

Very few longitudinal studies for bats investigate quantitative measures of energetically challenging portions of the annual cycle. In fact, this is the first study to apply commercially available hormone assays to multiple species of wild-caught pteropodid bats. While some previous studies have paired histology with hormonal assays to confirm reproductive stage classification and establish reference ranges for different phases of the reproductive cycle (Buchanan and Younglai 1986; Santiago *et al*. 2020; Beguelini *et al*. 2021), field studies focused on threatened species, including many pteropodids, cannot sacrifice whole animals. Our study successfully applied a mixture model approach to establish ranges for these phases corresponding to different hormones. However, because we did not sacrifice whole animals and collect histological data or detailed field notes on estimated embryos size, these hormone concentrations cannot be used to distinguish between early and late-stage pregnancies. Despite this, we showed that estradiol and progesterone measured in commercially available kits can be used as a tool to identify pregnancy status in the field – especially when paired with pre-existing knowledge of the seasonality of the reproductive calendar and supported by in-field abdominal palpation. The methods presented here serve as an important first step in applying quantitative non-destructive measures to determine pregnancy in bat field studies; our results present important implications for how to estimate population structure for conservation.

Overall, this is one of the first studies to understand the hormonal basis of reproductive cycling in Malagasy pteropodids. Moving forward, additional reliably collected, longitudinal data, and novel techniques (e.g., point-of-care devices, tracking, and/or recapturing) are needed to document and quantify various behaviors and life-stages, including foraging, gestation, and lactation. With the increase in technology available for field-based studies and remote-veterinary needs, there is an opportunity to obtain ultrasonic measures of multiple stages of pregnancy. Pairing measures of hormones with ultrasonic measures may be the only way to determine reference values for reproductive hormones in species which have unknown or unclear reproductive strategies or cycles without sacrificing the animal (though note, Moore *et al*. 2025). Accurate quantification of the Madagascar pteropodid bat reproductive calendar is an essential prerequisite to formulating quality demographic predictions to inform these species’ conservation, as well as elucidating the role that reproductive hormones play in modulating physiological changes in bat health.

## Data statement

All reproductive data, metadata, protocols, and statistical code used in all analyses is available for download from our public Github repository at: https://github.com/brooklabteam/Hormone-bats.git

## Authorship statement

ECR conducted the laboratory assays and data analysis, made the figures, and co-wrote the manuscript. GKD conducted laboratory validation and provided feedback for statistical analysis and writing. AA, SA, HCR, and CEB collected the longitudinal samples. FG assisted with writing and provided funding. CEB assisted with data analysis and figures, co-wrote the manuscript, and provided funding. All authors provided feedback on the final version of the manuscript.

## Acknowledgements

This research was funded by grants to ECR (Grainger Bioinformatics Center – Field Museum of Natural History) and CEB (Branco Weiss “Society in Science” fellowship; Walder Foundation ‘Biota’ Award, NIH DP2AI171120). We thank the Brook lab for comments on an earlier draft of this manuscript and DeeAnn Reeder for input on the hormonal ELISA methods. Sarah Guth, Anecia Gentles, and Fifi Ravelomanantsoa provided significant field support. The University of Antananarivo Mention de Zoologie et Biodiversité Animale aided in obtaining research and export permits. Finally, we also thank the Lubee Bat Conservancy (Gainesville, FL) for contributing sera to our validation process.

## Notes

**Declaration of competing interest:** The authors declare no conflict of interest.

https://github.com/brooklabteam/Hormone-bats

### Competing Interest Statement

The authors have declared no competing interest.

### Summary of Updates

We have re-analyzed the data and updated the results in response to reviewer feedback.

