## Appendix 1 for "Endocrinology reveals seasonal reproductive dynamics in three Malagasy fruit bats"

**ELISA validation results**

*Dilution linearity*

For all dilution linearity tests, we combined three to six plasma samples per species (n=6 samples *R. madagascariensis*; n=3 samples *P. rufus*; n=4 samples *E. dupreanum*) into species-specific plasma pools, then carried out 1:2, 1:4, 1:8, 1:16, 1:32, and 1:64 dilutions on each pool. We used a commercial ELISA kit (see main text) to assay the concentration of each initial species-specific pool and all downstream dilutions in the series.

For the dilution linearity series for progesterone, a linear regression evaluating the measured hormone concentration (y-axis) as predicted by dilution ratio (x-axis) for all three species recovered R^2^ values over 0.97 (Figure S1). For *R. madagascariensis*, normality assumptions were assessed using Shapiro-Wilk (W = 0.8560, p = 0.2144) and Kolmogorov-Smirnov (D = 0.2322, p > 0.10), and residual distributions were not significant (p> 0.05). For *P. rufus*, normality assumptions were assessed using Shapiro-Wilk (W = 0.8728, p = 0.2377) and Kolmogorov-Smirnov (D = 0.2335, p > 0.10), and residual distributions were not significant (p> 0.05). For *E. dupreanum*, normality assumptions were assessed using Shapiro-Wilk (W = 0.8610, p = 0.2317) and Kolmogorov-Smirnov (D = 0.2550, p > 0.10), and residual distributions were not significant (p> 0.05) (Figure S1). For progesterone, a Runs test showed no significant non-random residual structure for all three species (p > 0.05).

For the dilution linear series for estradiol, a linear regression evaluating the measured hormone concentration (y-axis) as predicted by dilution ratio (x-axis) for all three species recovered R^2^ values over 0.95 (Figure S1). For *R. madagascariensis*, normality assumptions were assessed using Shapiro-Wilk (W = 0.8620, p = 0.2355) and Kolmogorov-Smirnov (D = 0.2921, p > 0.10), and residual distributions were not significant (p> 0.05). For *P. rufus*, normality assumptions were assessed using Shapiro-Wilk (W = 0.7950, p = 0.0737) and Kolmogorov-Smirnov (D = 0.3370, p > 0.05), and residual distributions were not significant (p> 0.05). For *E. dupreanum*, normality assumptions were assessed using Shapiro-Wilk (W = 0.8166, p = 0.1099) and Kolmogorov-Smirnov (D = 0.2921, p > 0.10), and residual distributions were not significant (p> 0.05) (Figure S1). For estradiol, a Runs test showed no significant non-random residual structure for all three species (p > 0.05).

For the dilution linear series for testosterone, a linear regression evaluating the measured hormone concentration (y-axis) as predicted by dilution ratio (x-axis) for all four species (including an additional species *Rousettus aegyptiacus*) recovered R^2^ values over 0.97 (Figure S2). For *R. madagascariensis*, normality assumptions were assessed using Shapiro-Wilk (W = 0.7491, p = 0.0292) and Kolmogorov-Smirnov (D = 0.3810, p =0.0168), and residual distributions were not significant (p> 0.05). For *P. rufus*, normality assumptions were assessed using Shapiro-Wilk (W = 0.8641, p = 0.2035) and Kolmogorov-Smirnov (D = 0.3217, p > 0.05), and residual distributions were not significant (p> 0.05). For *E. dupreanum*, normality assumptions were assessed using Shapiro-Wilk (W = 0.8989, p = 0.4040) and Kolmogorov-Smirnov (D = 0.2531, p > 0.10), and residual distributions were not significant (p> 0.05) (Figure S2). For *R. aegyptiacus*, normality assumptions were assessed using Shapiro-Wilk (W = 0.9793, p = 0.9307) and Kolmogorov-Smirnov (D = 0.1512, p > 0.10), and residual distributions were not significant (p> 0.05) (Figure S2). For estradiol, a Runs test showed no significant non-random residual structure for all three species (p > 0.05).

*Parallelism*

For parallelism tests, we compared the slope of the dilution linearity curve for each bat species and hormone concentration against the standard curve for each assay. For progesterone, serially-diluted pooled serum samples from all three species were parallel to the standard curve, and a Runs test showed no significant non-random residual structure for all three species compared to the standard curve (p > 0.10) (Figure S3). For estradiol, serially-diluted pooled serum samples from all three species were parallel to the standard curve and a Runs test showed no significant non-random residual structure for all three species compared to the standard curve (p > 0.05) (Figure S3). For testosterone, serially-diluted pooled serum samples from all four species were parallel to the standard curve, and a Runs test showed no significant non-random residual structure for all three species compared to the standard curve (p > 0.05) (Figure S3).

*Spike and Recovery*

For progesterone, the average recovery percentage (%) was 104.16 ± 6.90% for *R. madagascariensis*, 93.93 ± 2.19% for *P. rufus*, and 97.86 ± 6.52% for *E. dupreanum* (Table S1; n=5 samples per species; mean ± standard error mean (SEM)). For estradiol, the average recovery % was 88.13 ± 2.28% for *R. madagascariensis*, 102.00 ± 6.86% for *P. rufus*, and 93.18 ± 1.99% for *E. dupreanum* (Table S1; n=5 samples per species; mean ± SEM). For testosterone, the average recovery % was 120.90 ± 3.93% for *R. madagascariensis*, 107.80 ± 8.45% for *P. rufus*, 146.30 ± 18.93% for *E. dupreanum*, and 107.80 ± 8.454% for *P. rufus*, 121.60 ± 8.16% for *R. aegyptiacus* (Table S1; n=5 samples per species; mean ± SEM). Acceptable spike recoveries should be between 80-120% (Selby et al., 2018).

**See attached excel spreadsheet.**

**Table S1.** Raw spike and recovery values for all three assay validations (progesterone, estradiol, and testosterone) for *R. madagascariensis, E. dupreanum*, and *P. rufus.* Values for *R. aegyptiacus* are only provided for testosterone.

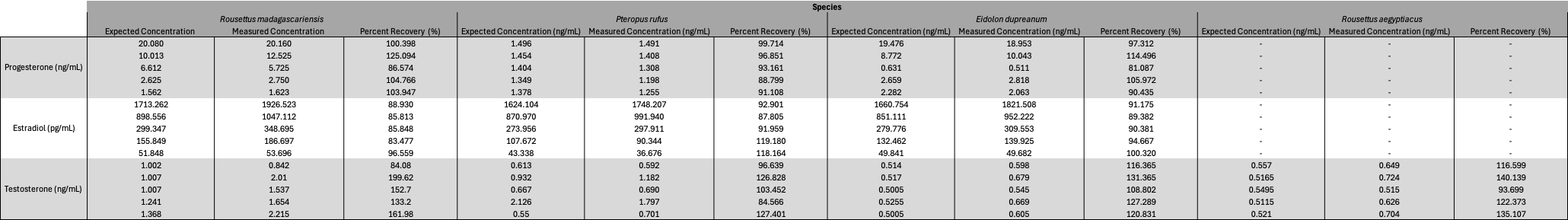

** see excel file for larger resolution.

**Table S2. Cutoff values for the selected species-hormone combinations.** Values are based on a cluster analysis approach outlined in the main manuscript methods.

| Bat species | Hormone | n clust | Id_clust | Prop | Mean | Sd | Cutoff_clust_lci | cutoff_clust | Cutoff_clust_uci |
| --- | --- | --- | --- | --- | --- | --- | --- | --- | --- |
| *Pteropus rufus* | PCA1 | 2 | 1 | 0.877 | 2.529 | 0.876 | 4.840 | **4.504** | 4.840 |
| *Pteropus rufus* | PCA1 | 2 | 2 | 0.123 | 4.888 | 0.286 |  |  |  |
| *Eidolon dupreanum* | PCA1 | 3 | 1 | 0.460 | 1.453 | 0.205 | 6.210 | **6.210** | 6.210 |
| *Eidolon dupreanum* | PCA1 | 3 | 2 | 0.491 | 2.638 | 0.813 |  |  |  |
| *Eidolon dupreanum* | PCA1 | 3 | 3 | 0.049 | 7.114 | 1.326 |  |  |  |
| *Rousettus madagascariensis* | PCA1 | 2 | 1 | 0.426 | 1.537 | 0.337 | 2.478 | **2.478** | 2.478 |
| *Rousettus madagascariensis* | PCA1 | 2 | 2 | 0.574 | 3.678 | 1.141 |  |  |  |

**Table S3. Average concentrations of each reproductive hormone by species and status.** Values represent the raw hormone concentrations as determined by commercially available assay. Status was determined in the field. For p-values from ANOVAs, estradiol significance is listed first, then progesterone.

| **Species** | **Sex** | **Status** | **Estradiol**  **(ng/mL) ± st.dev** | **Progesterone**  **(pg/mL) ± st.dev** | **Testosterone**  **(ng/mL) ± st.dev** | **p-value** |
| --- | --- | --- | --- | --- | --- | --- |
| ***E. dupreanum*** | Male | J |  |  | 50.95 ± 98.08 | p=0.058 |
|  |  | A |  |  | 176.38 ± 170.26 |  |
|  | Female | J | 94.95 ± 31.4 | 3.03 ± 1.46 |  | p<0.001; p<0.001 |
|  |  | NL | 91.81 ± 70.18 | 2.23 ± 1.68 |  |  |
|  |  | P | 393.23 ± 308.59 | 12.5 ± 6.98 |  |  |
|  |  | L | 84.73 ± 49.74 | 1.46 ± 0.85 |  |  |
| ***R. madagascariensis*** | Male | J |  |  | 5.76 ± n.a. | p=0.706 |
|  |  | A |  |  | 7.87 ± 5.53 |  |
|  | Female | J | 109.8 ± 48.24 | 8.06 ± 17.39 |  | p<0.001; p<0.001 |
|  |  | NL | 116.51 ± 43.28 | 9.2 ± 8.32 |  |  |
|  |  | P | 219.19 ± 132.8 | 26.16 ± 28.27 |  |  |
|  |  | L | 65.98 ± 49.45 | 3.84 ± 7.89 |  |  |
| ***P. rufus*** | Male | J |  |  | 16.57 ± 11.05 | p=0.031 |
|  |  | A |  |  | 132.45 ± 124.61 |  |
|  | Female | J | 26.84 ± 35.36 | 50.98 ± 68.76 |  | p=0.109; p=0.623 |
|  |  | NL | 155.25 ± 198.64 | 173.68 ± 298.38 |  |  |
|  |  | P | 863.1 ± 635.52 | 562.59 ± 455.12 |  |  |
|  |  | L | 49.82 ± 31.98 | 20.63 ± 18.96 |  |  |

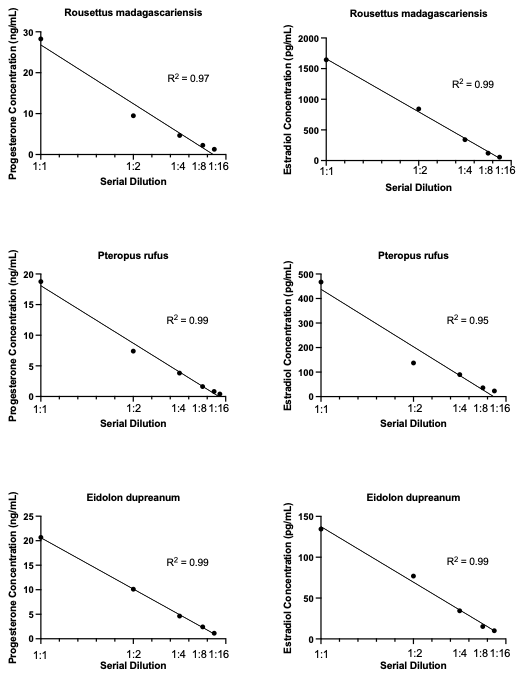

**Figure S1.** Linear relationships for quantification of reproductive markers in our three species of interest. Linear relationships between serial dilution and measured concentration for progesterone (left panels) and estradiol (right panels) in *Rousettus madagascariensis*, *Pteropus rufus*, and *Eidolon dupreanum* (top to bottom). Linear regression lines indicate assay performance and were used to validate hormone concentrations in biological samples.

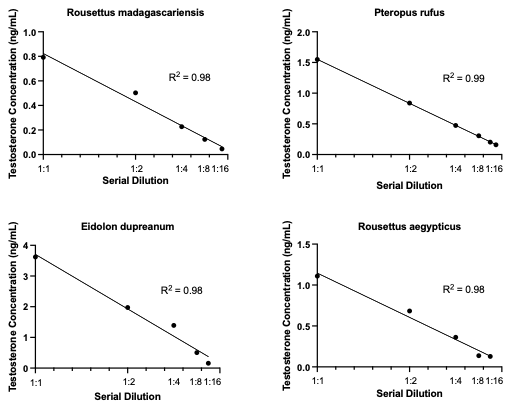

**Figure S2.** Linear relationships for quantification of reproductive markers in our three species of interest and *Rousettus aegyptiacus*. Linear relationships between serial dilution and measured concentration for testosterone in *Rousettus madagascariensis*, *Pteropus rufus*, *Eidolon dupreanum,* and *Rousettus aegyptiacus.* Linear regression lines indicate assay performance and were used to validate hormone concentrations in biological samples.

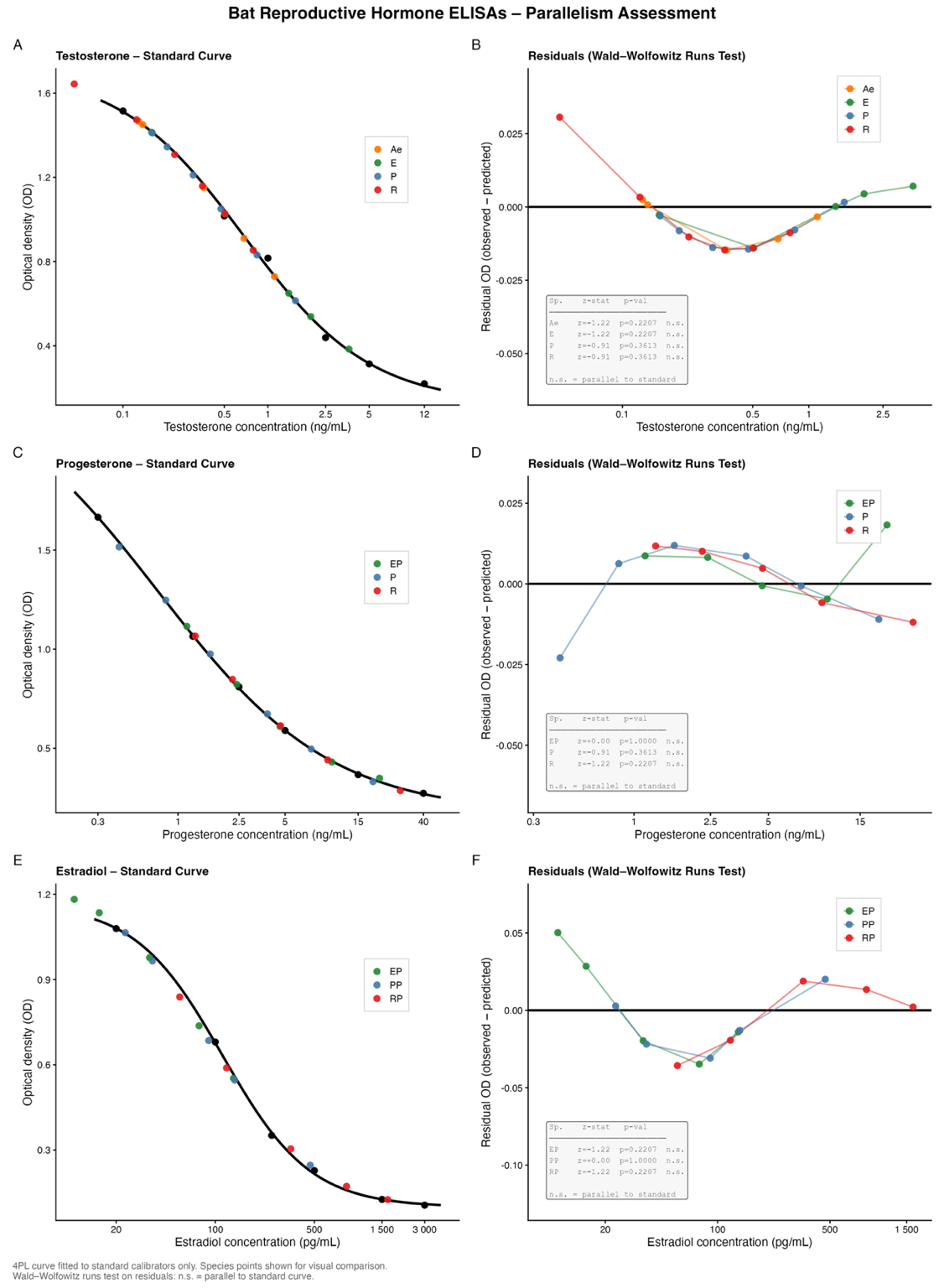

**Figure S3.** Standard curves and parallelism validation for testosterone, progesterone, and estradiol immunoassays. (A, C, E) Four-parameter logistic standard curves (black line and points) for testosterone (ng/mL), progesterone (ng/mL), and estradiol (pg/mL), with extracted sample dilutions overlaid (colored points). (B, D, F) Residuals (observed − predicted OD) from Wald–Wolfowitz runs tests assessing parallelism between serial dilutions of each sample type and the standard curve. All sample types were parallel to the standard (n.s.; see inset tables for z-statistics and p-values), validating the use of these assays for quantifying hormone concentrations in the sample matrices tested. Abbreviations: Ae = *R. aegyptiacus*, E = *E. dupreanum*, P = *P. rufus*, R = *R. madagascariensis*.

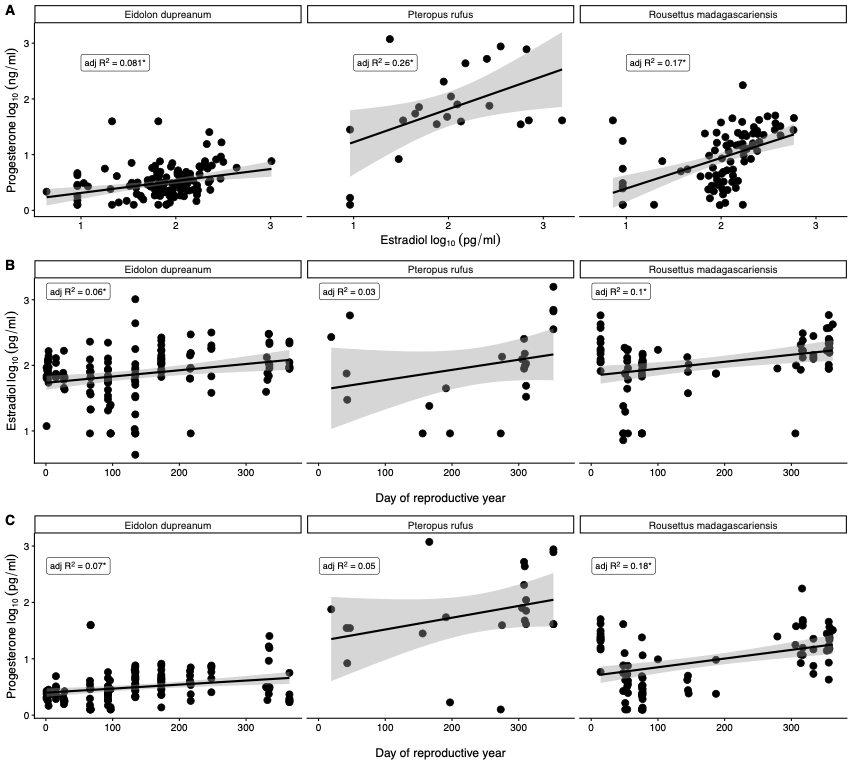

**Figure S4.** (A) Correlation between estradiol and progesterone in all three species of fruit bats. Correlation of estradiol (B) and progesterone (C) and reproductive day of the year. The significant correlation of estradiol and progesterone led us to conduct a PCA on the two hormones combined with day of reproductive year. This approach is described further in the main text methods. Adjusted R^2^ is noted on each plot, the presence of an asterisks indicates significance at p<0.05/

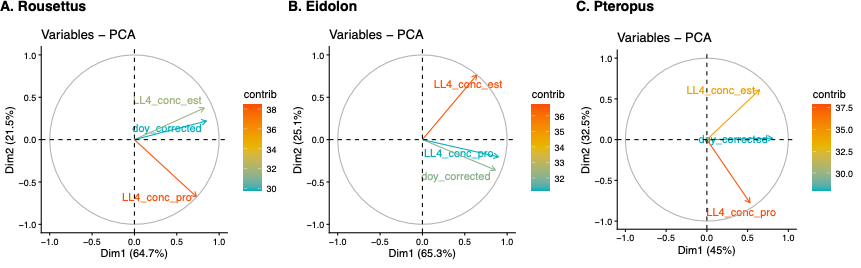

**Figure S5. Principal component analysis (PCA) biplots of reproductive hormone variables for three fruit bat species**: **Rousettus madagascariensis (A), Eidolon dupreanum (B), and Pteropus *rufus* (C).** Arrows indicate the contribution (contrib) of each variable — estradiol concentration (LL4_conc_est), progesterone concentration (LL4_conc_pro), and day of reproductive year (doy_corrected; day 0 corresponds to birth date) — to the first two principal components. Dim1 explains 64.7%, 65.3%, and 45.0% of variance in Rousettus, Eidolon, and Pteropus, respectively; Dim2 explains an additional 21.5%, 25.1%, and 32.5%. Arrow color reflects each variable's relative contribution to the PCA, with warmer colors indicating greater contribution.

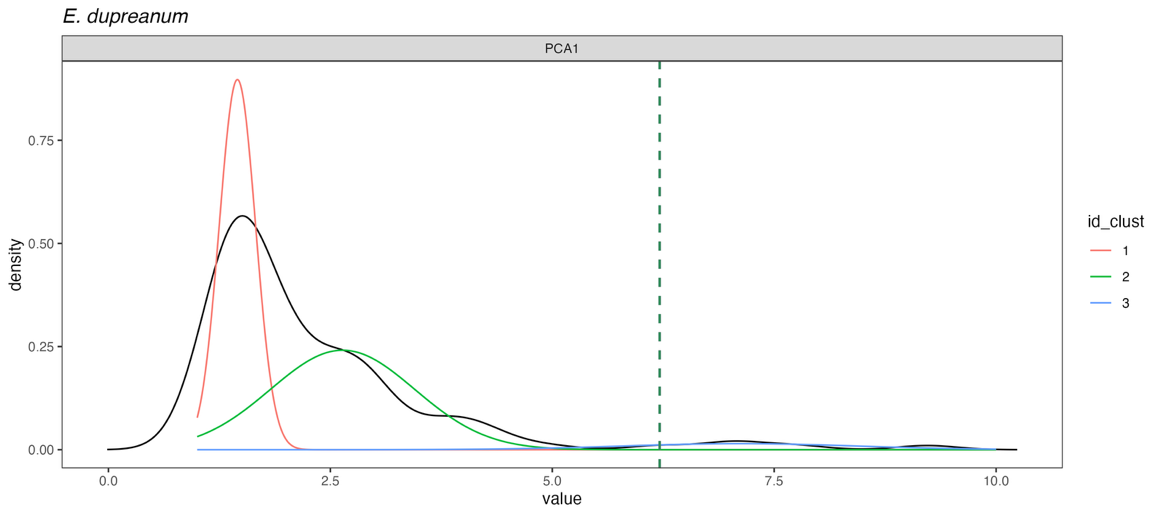

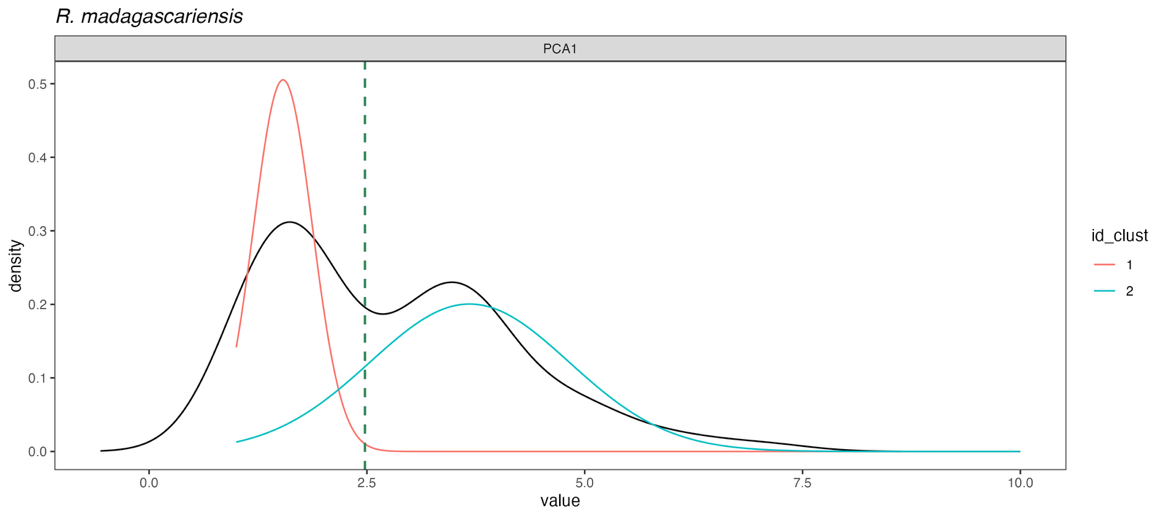

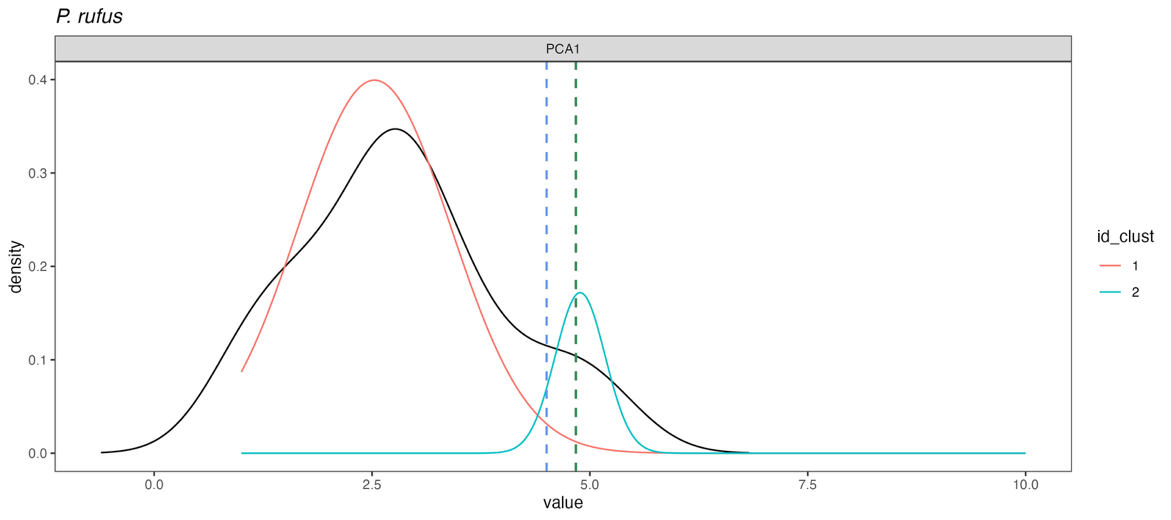

**Figure S6.** Distribution of PCA values and location of lower confidence interval (blue dashed vertical line– 60%), cutoff as determined by 70% confidence (red dashed vertical line), and upper confidence interval (green dashed vertical line – 85%). Methods for determining cutoffs are presented in the main text methods. In the case of *P. rufus* and *R. madagascariensis*, the lci (60%) and 70% confidence interval overlapped and thus plots only show two lines.

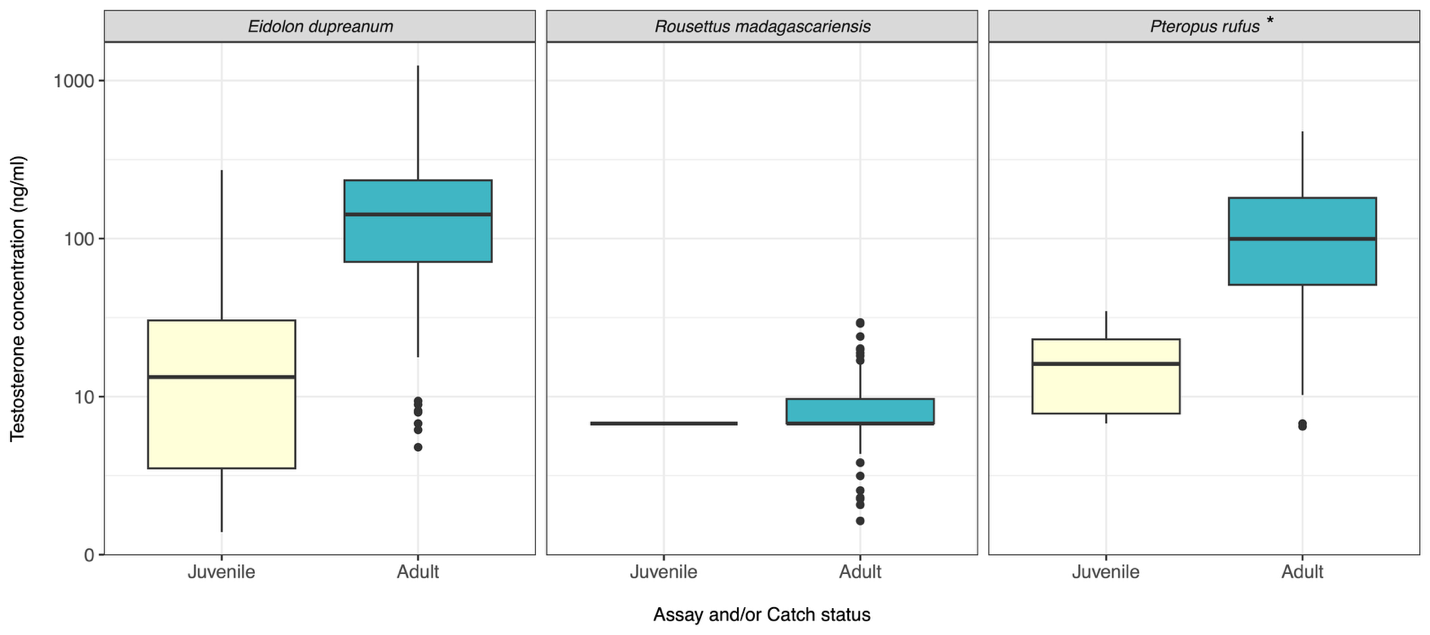

**Figure S7.** Testosterone (ng/ml) for male bats, plotted by reproductive status. Adult *P. rufus* were the only species where adults had significantly higher testosterone than juvenile individuals. Asterisks in the header indicate a significant (p<0.05) ANOVA result.

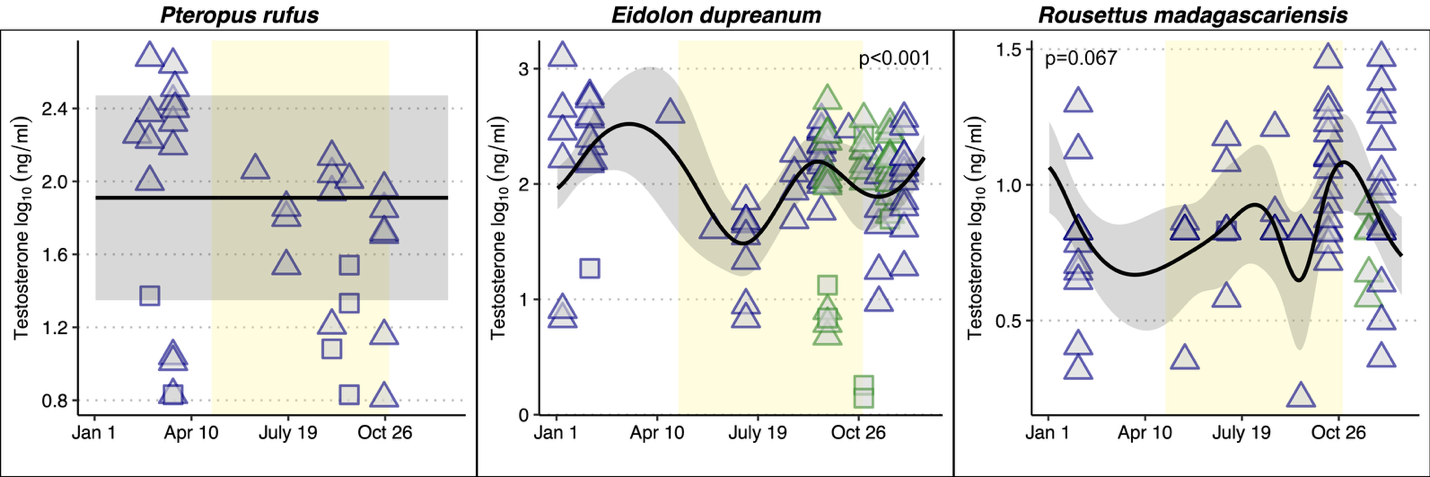

**Figure S8.** Seasonal patterns of testosterone for all three species of bats. *P. rufus* on left, *E. dupreanum* in the middle, and *R. madagascariensis* on the right. Shape of the points are indicated by catching status – triangles are adults, whereas squares are juveniles. Points outlined in dark blue are from the District of Moramanga whereas points outlined in green are from Ankarana. Solid line indicates generalized additive model trends with 95% confidence intervals shaded in gray (GAMs were fit to District of Moramanga data only). Background yellow ribbon corresponds to the winter season of nutritional deficit in the District of Moramanga sites.
